# End-to-end plaque counting and virus titration from laboratory plate images with deep learning

**DOI:** 10.64898/2026.09.04.749375

**Authors:** Eugenia Moris, Alicia Costábile, Sebastián Rey, Irene Ferreiro, Joaquín Hurtado, Lizandra Lissette Luciano, Matías Villagrán, Aisha Espino Vázquez, Jomari Ramos, Isadora Monteiro, María Victoria de Santiago, Pilar Moreno, Gonzalo Moratorio, José Ignacio Orlando

## Abstract

Plaque assays are the gold standard for quantifying infectious virus, yet plaque enumeration is still routinely performed manually, making virus titration labor-intensive, subjective, and difficult to standardize across analysts and laboratories. Existing automated methods primarily address individual tasks, such as plaque segmentation or counting, but do not provide an integrated workflow from plate images to biological quantification. In this paper we present Titra, an end-to-end workflow for automated cytopathic effect (CPE)-based virus titration from standard photographs of plaque assay plates. The workflow combines automatic well detection, plaque segmentation, post-processing for instance separation, plaque counting, and plaque-forming units per millilitre (PFU/mL) estimation within a single web-based platform that also enables experiment management and expert review. The approach was evaluated using images from three viral species (Mayaro virus, Coxsackievirus B3, and vaccinia virus), two plate formats (6- and 12-well), and heterogeneous image acquisition conditions, including both a newly curated dataset and the public VACVPlaque dataset. Automated plaque counts showed strong agreement with manual annotations (Pearson correlation coefficients of 0.98 for MAYV/CVB3 and 0.88 for VACV), while PFU/mL estimates closely matched manual calculations for the MAYV/CVB3 dataset (Pearson r=0.975). Comparative experiments against U-Net, StarDist, HSD-WBR, and PyPlaque demonstrated competitive segmentation and counting performance, with the proposed approach achieving the highest Dice and mAP on the public VACVPlaque dataset.

**Author summary:** Plaque assays are widely used to measure infectious virus, but manually counting plaques is labor-intensive and can vary between researchers. We developed Titra, an automated workflow that analyzes photographs of plaque assay plates, detects assay wells, counts viral plaques, and calculates virus concentration in plaque-forming units per milliliter (PFU/mL). The workflow also provides a web-based interface that allows researchers to review and correct automated results when needed. We evaluated the approach using images of Mayaro virus, Coxsackievirus B3, and vaccinia virus acquired from different plate formats and imaging conditions. Automated plaque counts showed strong agreement with manual annotations and with counts from four independent experts. Although performance decreased in wells containing highly confluent plaques, these cases were also the most challenging for manual annotation. Our findings demonstrate that automated plaque assay analysis can substantially reduce routine manual effort while preserving expert review and providing reliable virus titration.

## Introduction

Plaque assays remain the gold-standard method for quantifying infectious virus, but plaque enumeration is still commonly performed by manual visual inspection, making the process labor-intensive, subjective, and difficult to standardize across analysts and laboratories [1]. To reduce this problem, several software tools have been proposed to automate specific stages of plaque assay analysis. Classical image-processing approaches, including Plaque2.0 [2], Viridot [3], ViralPlaque [4], the Plaque Size Tool [5], and PyPlaque [6], provide automated plaque detection or counting, but typically rely on user-defined parameters, manual well selection, or are tailored to specific assay formats. More recently, deep-learning approaches have been applied to this task, including general-purpose segmentation architectures such as U-Net [7] and StarDist [8]. In a more specialized manner, HSD-WBR [9, 10] combines Hydra and StarDist for plaque detection on bright-field 6-well plates. While these methods have substantially improved automated plaque segmentation, they primarily focus on isolated tasks, such as plaque segmentation or counting, rather than providing a complete workflow for routine virus titration.

Recent advances in foundation models for image segmentation provide an opportunity to improve those results. The Segment Anything Model (SAM) demonstrated strong generalization across diverse imaging domains while requiring relatively little task-specific annotation [11]. At the same time, publicly available datasets such as VACVPlaque [12] have enabled the evaluation of automated methods on realistic, label-free plaque assay images acquired using standard laboratory photography. Together, these developments make it feasible to design robust workflows that operate under heterogeneous imaging conditions without requiring specialized acquisition hardware. However, existing approaches still generally terminate at plaque segmentation or counting, leaving subsequent steps–including well localization, PFU/mL calculation, quality control, and experiment management–to manual analysis.

In this study, we present Titra (https://titra.app/), an end-to-end workflow for computer-aided CPE-based virus titration directly from plaque assay plate photographs. The proposed workflow automatically localizes assay wells, segments and counts plaques, computes plaque-forming units per millilitre (PFU/mL), and integrates these analyses into a web-based platform that allows expert review and experiment management. Unlike previous methods that automate individual components of plaque assay analysis, Titra provides a complete workflow from raw plate images to biological quantification. We evaluate the proposed approach on three viral species (Mayaro virus, Coxsackievirus B3, and vaccinia virus), two plate formats, and heterogeneous acquisition conditions, comparing its segmentation performance with state-of-the-art methods and validating its plaque counts and PFU estimates against expert annotations. In addition, we introduce the MAYV/CVB3 dataset, collected at the Institut Pasteur from Montevideo (Uruguay) under routine laboratory conditions using smartphone photography, which will be publicly released upon acceptance of this manuscript jointly with our code implementation.

## Materials and methods

### Proposed approach

The complete workflow is illustrated in Fig. 1. A plaque assay plate image acquired using a standard camera is first uploaded to Titra together with the corresponding experimental metadata. The image is then processed by a SAM2-based [13] well-segmentation model, which identifies individual wells and determines their positions within the plate. Each detected well is subsequently analyzed by the SAM-based plaque-segmentation model, followed by the post-processing stage described below to separate overlapping plaques and estimate plaque counts. We adopted this foundation-model approach, using SAM2 [13] for well segmentation and the Segment Anything Model (SAM) [11] for plaque segmentation, because plaque assay images exhibit substantial variability across viral species, plate formats, and image acquisition conditions, and these models have been observed to generalize well beyond to multiple tasks. In particular, SAM2 extends the promptable segmentation paradigm introduced by SAM and has demonstrated strong accuracy and efficiency across diverse image and video segmentation benchmarks [13]. Similarly, SAM-based models have shown excellent transferability to biomedical imaging applications, including microscopy (*µ*SAM) [14] and universal medical image segmentation (MedSAM) [15]. This modular design also enables future replacement or fine-tuning of individual components without modifying the overall workflow.

**Fig 1.**
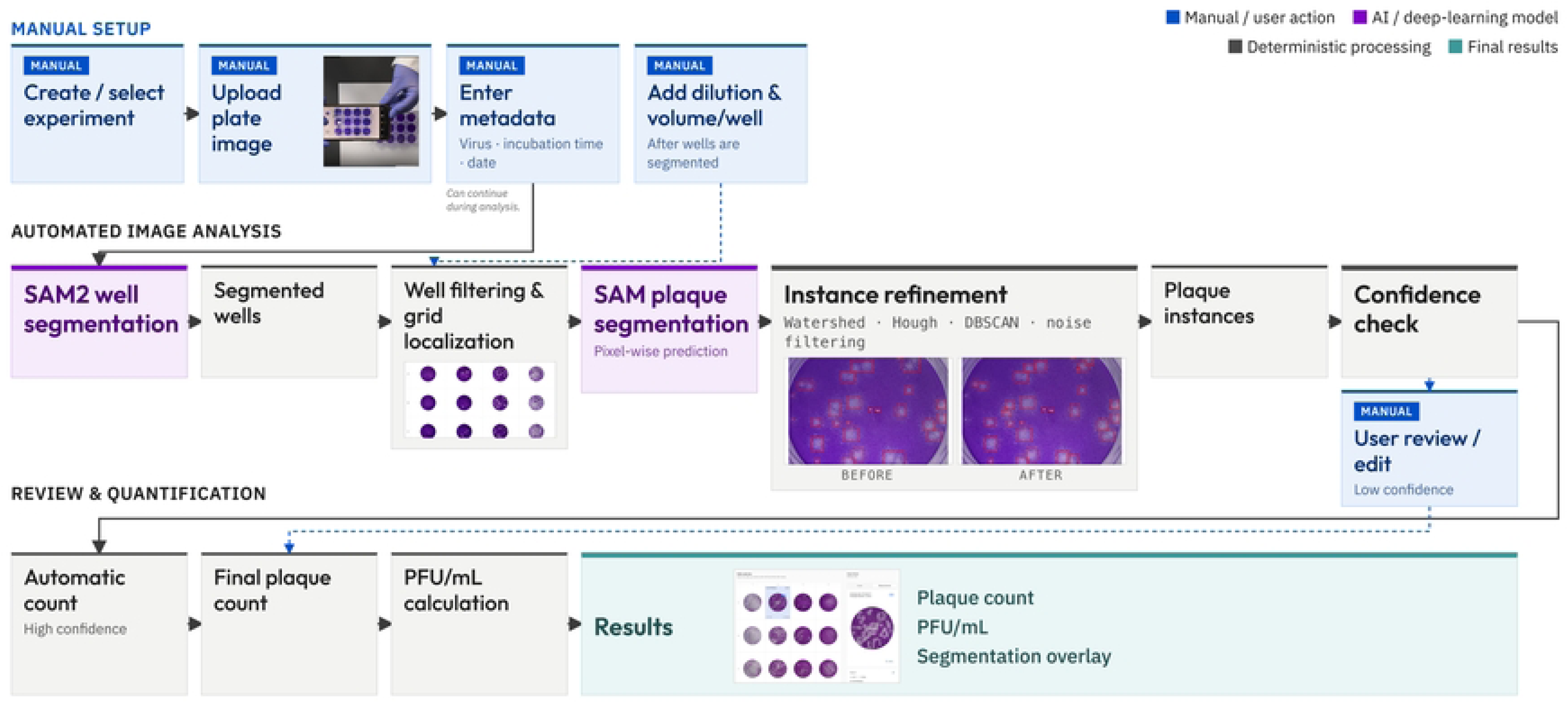
End-to-end workflow for automated plaque assay quantification in *Titra*. A full-plate image is processed using a SAM2-based well-segmentation module to identify and localize individual wells (Well Detection module). Each segmented well is subsequently analysed by a SAM-based plaque-segmentation model, followed by post-processing to resolve overlapping plaques (Plaque Detection module). The resulting plaque counts are combined with dilution and inoculation information to compute PFU/mL values.

While plaque segmentation is performed automatically, users can simultaneously review or update the associated experimental metadata. To this end, right after plaque detection, the workflow presents the segmentation results through the Titra interface, allowing users to inspect and, if necessary, manually correct individual plaque detections before confirming the final counts. Once the review is completed, the validated plaque counts are automatically combined with the corresponding dilution and inoculation-volume information to calculate plaque-forming units per millilitre (PFU/mL), providing the final virus titration result.

### Well segmentation

Well segmentation is performed in a zero-shot setting using SAM2 initialised from the pre-trained checkpoint `sam2.1_hiera_base_plus.pt` with configuration file `sam2.1_hiera_b+.yaml`. This variant was chosen to balance segmentation capacity and computational efficiency, with no fine-tuning applied.

Prior to inference, input images are rescaled with preserved aspect ratio, setting the longest side to 256 pixels. Candidate masks are generated using the SAM2 automatic mask generator (`points_per_side=12, min_mask_region_area=100`). The built-in post-processing is disabled in favor of our custom filtering stage.

Candidate masks are then filtered using geometric constraints to remove spurious detections and non-well regions. For each candidate region, we compute the area, perimeter, circularity (*C* = 4*πA/P*^2^), and eccentricity. Only regions exceeding 100 pixels in area and exhibiting circularity greater than 0.7 are retained. For each validated mask, a bounding box is computed and mapped back to the original image resolution.

To determine the plate layout, the validated well masks are further processed to estimate the relative position of each well within the grid. This positional information is used to assign each well to its corresponding row and column, enabling correct association with the experimental dilution scheme during subsequent PFU/mL calculation.

This design prioritises robust and reliable well detection under CPU-only inference, ensuring scalability and compatibility with standard laboratory computing environments.

### Plaque segmentation

Plaque segmentation is performed using the image encoder of SAM [11], which is used as a frozen feature extractor owing to its strong generalisation capabilities and its ability to capture fine-grained object boundaries. In particular, we used the `ViT-B` Vision Transformer backbone, keeping the encoder weights frozen throughout training to preserve the pre-trained visual representations learned from large-scale image data. This strategy—freezing a large pre-trained encoder and training only a task-specific decoder—has been explored successfully in prior studies [16, 17].

A lightweight convolutional decoder was attached to the frozen encoder to predict plaque segmentation masks. The decoder consists of a 3 × 3 convolutional layer followed by a ReLU activation and a final 1 × 1 convolution producing a single-channel output. Only the decoder parameters were optimised, enabling efficient training with fewer computational resources while retaining the representational capacity of the SAM encoder. The model outputs pixel-wise plaque probability maps.

Model training was implemented using the PyTorch Lightning framework [18] to support reproducible and structured experimentation. Optimisation was performed using Adam optimiser [19] and a binary cross entropy loss [20] objective, with learning rates explored in the range 1 × 10*^−^*^5^ to 1 × 10^−3^. Training was conducted for up to 50 epochs with a batch size of 2 on a NVIDIA GeForce RTX 4090. Performance was monitored using intersection-over-union (IoU), Dice coefficient, and pixel accuracy on both training and validation sets. For multi-dataset training (e.g., MAYV/CVB3 and VACV), balanced mini-batch sampling was applied to mitigate dataset imbalance. Each batch contained 50% randomly selected samples from each dataset. When dataset sizes differed, the number of batches per epoch was limited by the smaller dataset.

Input data consisted of 1024 × 1024 RGB images corresponding to individual wells extracted from multi-well assay plates, paired with binary GT masks. Images were resized to the target resolution and standardised using ImageNet statistics, while GT masks were processed to ensure binary consistency. To improve generalisation across viruses and experimental conditions, we applied extensive data augmentation using a RandAugment-inspired pipeline [21, 22]. The number of augmentation operations per sample (`num_ops` ∈ {2, 4, 6, 8}) and the augmentation magnitude (`magnitude` ∈ {3, 7, 9, 11}) were selected via hyperparameter exploration. Geometric transformations (horizontal and vertical flips, and random rotations) were applied synchronously to images and masks, whereas colour-based augmentations were applied to images only and included `AutoContrast, Equalize, Solarize, ColorJitter, Sharpness`, and `Posterize`, as implemented in Torchvision [23].

### Plaque post-processing

Because the model produces pixel-wise probability maps without explicitly separating individual instances, a post-processing stage is required to distinguish touching plaques and successfully derive plaque counts. The proposed strategy combines morphological separation, geometric validation, and density-based filtering.

First, watershed segmentation is applied to the distance transform of the predicted binary mask. Local maxima were used as seed points to separate touching plaques. Hough circle detection was then used to identify approximately circular structures, validate plaque geometry, and recover potentially merged regions. A circularity threshold of 0.5 is applied to flag irregular or ambiguous candidates. A DBSCAN clustering procedure [24] is subsequently applied to merge nearby peaks and reduce over-segmentation artefacts in dense plaque configurations. Finally, a bounding box was generated for each detected instance, and detections smaller than 10 × 10 pixels are considered noise and discarded. Together, these morphological, geometric, and density-based refinements improve plaque delineation in crowded regions containing overlapping plaques. For comparison, we also evaluated plaque counts obtained using PyPlaque [25], a library that detects and counts plaques from segmentation masks generated from fluorescence or crystal-violet plaque assay images.

## Materials

### MAYV/CVB3 dataset

The MAYV/CVB3 dataset was developed at the Instituto Pasteur from Montevideo (Uruguay) and comprises 101 plaque assay images for training and validation, and 17 images for testing. All images were captured using a smartphone camera under ambient laboratory lighting, with the plates placed against a uniform white background to enhance contrast.

For plaque assays, Vero cells (ATCC CCL-81) were seeded in 12- and 6-well plates one day before infection at 2 × 10^5^ cells per well. Mayaro virus and Coxsackievirus B3 stocks were serially diluted (ten-fold) in serum-free DMEM. Cells were washed twice with PBS and infected with 100 *µ*L of each dilution for 1h at 37*^◦^*C. A semi-solid overlay (DMEM supplemented with 2% fetal bovine serum and 0.8% agarose; Invitrogen) was then added. After 45h at 37*^◦^*C, cells were fixed with 4% paraformaldehyde for 40min. The overlay was removed, wells were stained with 0.2% crystal violet for 40min, rinsed with tap water, air-dried, and imaged for plaque detection.

Manual well and plaque segmentations were created using the open-source annotation tool Label Studio [26]. During dataset curation, a small subset of images was excluded from the training and validation sets due to excessive plaque density or because presented fully transparent wells, which made accurate manual annotation infeasible. This filtering step aimed to ensure high-quality annotations and to prevent the inclusion of ambiguous cases that could introduce noise or bias during model training. Nonetheless, images containing partially visible or sparsely distributed plaques were retained to preserve dataset diversity and improve model generalisation. To avoid data leakage, images from the same experimental plate were kept within a single subset (either training, validation, or test).

### Vaccinia dataset

The vaccinia virus (VACV) dataset is a publicly available resource introduced by De *et al.* [12]. It comprises 211 digital photographs of 6-well tissue culture plates used for plaque assays. Each image includes manually annotated masks for both wells and plaques. The dataset was divided into training, validation, and test subsets containing 149, 43, and 22 images, respectively. The assays were performed on BSC40 cell monolayers infected with the VACV Western Reserve strain under various experimental conditions. Images were acquired using digital cameras. Given the availability of high-quality manual annotations, we used this dataset both for direct model evaluation and to assess the generalisation ability of our approach across datasets with differing imaging conditions and biological setups.

## Results

### Well detection results

The proposed well-detection model achieved highly accurate localization across both the MAYV/CVB3 and VACV datasets, maintaining high precision and recall over a broad range of IoU thresholds (Fig. 2).

**Fig 2.**
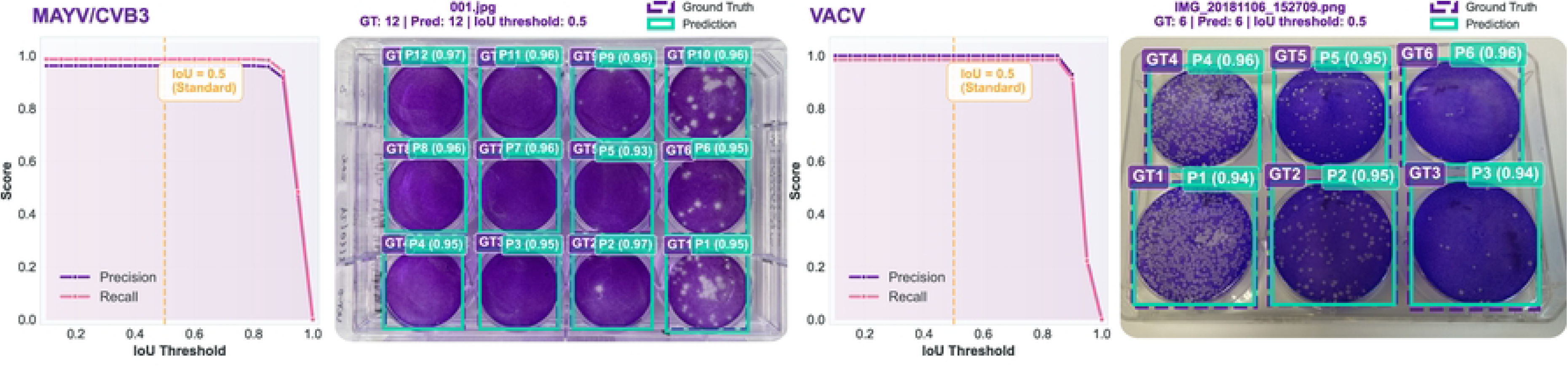
Automated well detection results in MAYV/CVB3 (left) and VACV (right). Left: precision and recall variations for different Intersection over Union (IoU) thresholds. Right: Representative examples of well detections, including ground truth (GT) boxes (purple) and predicted wells (light green).

Performance remained stable up to approximately an IoU threshold of 0.85, after which both precision and recall decreased as increasingly strict overlap criteria were applied. At the selected IoU threshold of 0.5, recall and precision exceeded 0.96 for both datasets, demonstrating reliable well localization despite differences in plate format and image acquisition conditions.

Representative examples (Fig. 2) illustrate accurate well localization across both evaluated datasets despite differences in plate format, camera viewpoint, and illumination conditions. In both examples, all wells were correctly detected with close agreement between the predicted masks and the manual annotations, demonstrating that the model remained robust to the heterogeneous acquisition conditions present in the datasets.

Failure cases (Fig. 3) corresponded to scenarios in which the contrast against the surrounding plate background was too low for the segmentation model to distinguish a boundary — i.e., the well’s appearance converged toward that of the background — most commonly (58% of cases) because heavy plaque formation had cleared the monolayer to near-background transparency, but also (42% of cases) from generally faint/low-contrast staining that kept the well dark yet indistinct.

**Fig 3.**
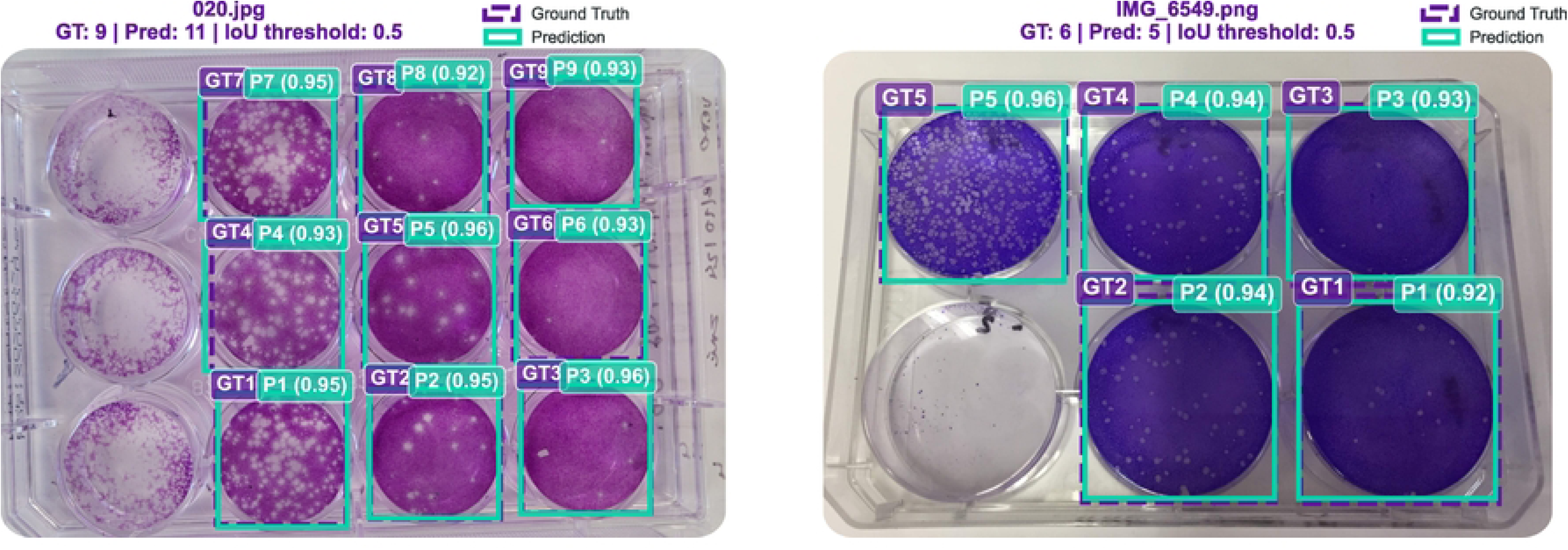
Examples of well-detection failure cases. Left: MAYV/CVB3 plate example where three out of twelve wells were not detected. Right: VACV plate example where one of six wells was missed.

### Plaque detection results

Plaque-counting performance differed across methods and datasets (Table 1). On MAYV/CVB3, U-Net achieved the lowest counting error (MAE: 1.81 ± 3.84 plaques per well), followed by Titra (3.45 ± 4.33). The difference between U-Net and Titra was statistically significant after Holm correction (*p <* 0.001), with Titra showing significantly higher counting error than U-Net on this dataset. HSD-WBR and StarDist showed higher errors (7.82 ± 12.79 and 10.43 ± 13.99, respectively), both significantly higher than Titra. On VACV, Titra achieved the lowest counting error (44.72 ± 112.41), significantly outperforming all three competing methods. Thus, U-Net provided the most accurate plaque counts on MAYV/CVB3, whereas Titra provided the most accurate counts on VACV.

**Table 1.** Comparison of plaque segmentation and plaque-counting performance across the MAYV/CVB3 and VACV datasets. Values are reported as mean ± standard deviation. Higher Dice, mAP, and F1 values indicate better segmentation/detection performance, whereas lower MAE values indicate more accurate plaque counting. Statistical significance for segmentation and counting comparisons is indicated using the symbols defined below the table.

|  | MAYV/CVB3 |  |  |  | VACV |  |  |  |
| --- | --- | --- | --- | --- | --- | --- | --- | --- |
| | Dice ↑ | mAP ↑ | F1 ↑ | MAE ↓ | Dice ↑ | mAP ↑ | F1 ↑ | MAE ↓ |
| U-Net [7] | **0.769 ± 0.28** | 0.111 ± 0.11 | **0.661 ± 0.34** | **1.81 ± 3.84<sup>‡</sup>** | 0.410 ± 0.30 | 0.018 ± 0.02 | 0.276 ± 0.32 | 85.11 ± 164.21<sup>γ</sup> |
| StarDist [8] | 0.622 ± 0.36 | **0.406 ± 0.41** | 0.539 ± 0.38 | 10.43 ± 13.99<sup>γ</sup> | 0.287 ± 0.26 | 0.098 ± 0.15 | 0.164 ± 0.20 | 80.41 ± 127.19<sup>γ</sup> |
| HSD-WBR [9, 10] | 0.504 ± 0.34 | 0.280 ± 0.35 | 0.429 ± 0.36 | 7.82 ± 12.79<sup>β†</sup> | 0.548 ± 0.27 | 0.184 ± 0.16 | 0.461 ± 0.30 | 46.11 ± 84.87<sup>α†</sup> |
| Titra (ours) | 0.423 ± 0.37 | 0.224 ± 0.31 | 0.397 ± 0.37 | 3.45 ± 4.33 | **0.649 ± 0.27\*** | **0.324 ± 0.25\*** | **0.564 ± 0.30** | **44.72 ± 112.41** |

Asterisks denote statistically significant improvements in segmentation performance over the competing methods ( $p < 0.05$ ). <sup>α</sup>, <sup>β</sup>, and <sup>γ</sup> indicate significantly higher plaque-counting errors than Titra at Holm-adjusted $p < 0.05$ , $p < 0.01$ , and $p < 0.001$ , respectively. <sup>‡</sup> indicates a significantly lower plaque-counting error than Titra ( $p < 0.001$ ). <sup>†</sup>For HSD-WBR, plaque counts were obtained using the post-processing procedure provided in the HSD-WBR code.

Segmentation and instance-detection metrics did not always follow the same ranking as counting performance. On MAYV/CVB3, U-Net achieved the highest Dice (0.769 ± 0.28) and detection F1 (0.661 ± 0.34), while StarDist achieved the highest mAP (0.406 ± 0.41). Although both StarDist and HSD-WBR achieved higher Dice and detection F1 than Titra, they produced larger counting errors. On

VACV, the results were more consistent across metrics: Titra achieved the highest Dice (0.649 ± 0.27), mAP (0.324 ± 0.25), and detection F1 (0.564 ± 0.30), together with the lowest counting MAE. The improvements in Dice and mAP were statistically significant.

To further characterize the different rankings observed for detection F1 and counting MAE on MAYV/CVB3, we analyzed errors in empty and non-empty wells separately. In wells containing plaques, Titra achieved a detection F1 of approximately 0.49–0.50, comparable to U-Net (∼ 0.52) and higher than StarDist (∼ 0.40–0.41). The main difference emerged in empty wells. Titra produced at least one false-positive plaque in 73% of empty wells, compared with 29% for StarDist, 16% for U-Net, and 9% for HSD-WBR. However, Titra’s errors were typically small, with approximately two spurious plaques per affected well. In contrast, StarDist produced false positives less frequently but occasionally generated much larger overcounts, reaching 43 spurious detections in a single well. Seven of its ten largest counting errors occurred in empty or near-empty wells, and these ten wells accounted for 35% of its total absolute counting error.

Representative plaque-segmentation examples are shown in Fig. 4(a). Across both datasets, isolated plaques were generally segmented correctly, whereas most discrepancies occurred in regions containing overlapping or densely packed plaques. Yellow arrows highlight representative missed or merged plaques that contributed to undercounting, while light-green arrows indicate representative spurious plaque detections corresponding to small plaque-like structures or image artefacts that were segmented as plaques despite being excluded from the manual annotations. These failure modes became increasingly frequent as plaque density increased, particularly in the VACV dataset, where extensive plaque overlap resulted in substantial undercounting. The qualitative examples also illustrate the marked differences between datasets, with VACV exhibiting substantially smaller and more densely distributed plaques than MAYV/CVB3.

**Fig 4.**
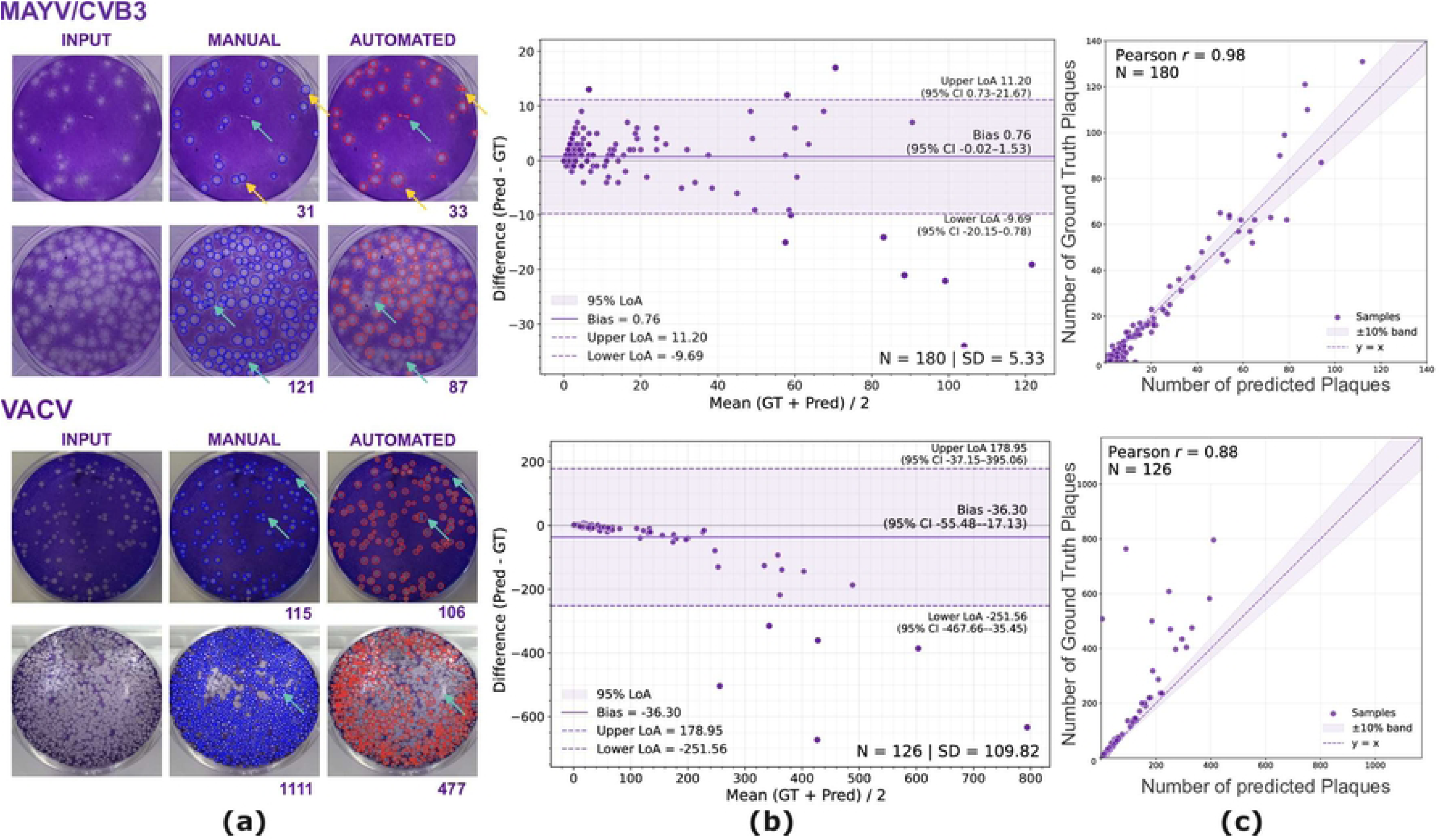
Automated plaque detection results in the MAYV/CVB3 and VACV datasets. (a) Representative plaque assay wells from the MAYV/CVB3 (top) and VACV (bottom) datasets–left to right: input images, manual annotations, and automated segmentation outputs, including plaque counts below; top to bottom: wells with low and high plaque density. (b) Bland–Altman plots showing agreement between automated predictions (Pred) and manual ground truth counts (GT) across all test wells. (c) Correlation plots comparing automated and manual plaque counts, with dashed lines denoting ±10% deviation bands.

Scatter plots comparing automated and manual plaque counts (Fig. 4(c)) demonstrated strong linear agreement across both datasets, with Pearson correlation coefficients of 0.98 for MAYV/CVB3 (*N* = 180) and 0.88 for VACV (*N* = 126). Although correlation remained high, the lower coefficient observed for VACV reflects the greater challenge posed by highly confluent plaques.

Bland–Altman analysis (Fig. 4(b)) further demonstrated good agreement between automated and manual plaque counts. For MAYV/CVB3, the mean difference (bias) was 0.76 plaques, with narrow limits of agreement (11.20 and −9.69). For VACV, the mean bias was −36.30 plaques, with wider limits of agreement (178.95 and −251.56), reflecting the substantially larger plaque counts present in this dataset. In both datasets, most observations remained within the limits of agreement. However, for VACV the differences became progressively more negative as plaque density increased, indicating systematic undercounting in highly confluent wells.

### Post-processing comparison with PyPlaque

Figure 5 illustrates the results obtained using the proposed post-processing stage for separating overlapping plaques after segmentation. Starting from the predicted segmentation mask, watershed-based instance separation identifies merged plaques and refines them into individual detections. The highlighted examples show representative regions in which a single connected segmentation is successfully separated into multiple plaque instances.

**Fig 5.**
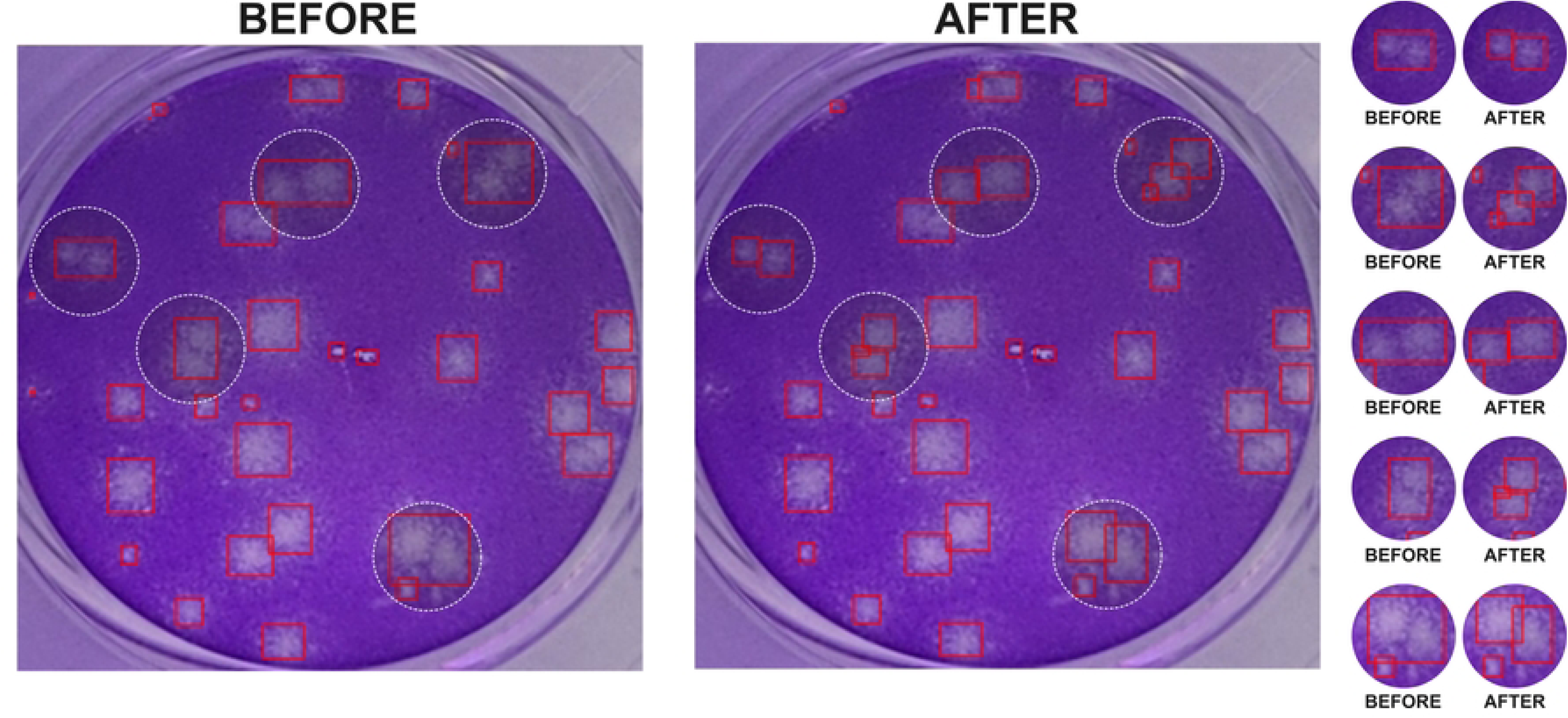
Qualitative results of our post-processing refinement for separating overlapping plaques in a representative well. Red bounding boxes indicate detected plaques, and white dashed circles highlight regions in which overlapping or merged plaques were identified and separated. The right panel presents magnified examples before and after separation into distinct detections.

To evaluate the contribution of the proposed post-processing independently of the segmentation model, we compared our watershed-based strategy with the connected-component counting approach implemented in PyPlaque [25]. Both methods were applied to the same predicted segmentation masks, ensuring that the post-processing stage was the only variable. Performance was evaluated using mean absolute error (MAE) for plaque counts and instance-level F1 score obtained through greedy IoU matching (IoU threshold = 0.5) between predicted and ground-truth plaque instances. Statistical significance was assessed using paired Wilcoxon signed-rank tests.

Across all evaluated wells (306 total), the proposed post-processing achieved a significantly higher instance-level F1 score than PyPlaque (0.481 vs. 0.424). The largest improvement was observed on the VACV dataset, where F1 increased from 0.542 to 0.612 (*p <* 0.001). On the MAYV/CVB3 dataset, the proposed method also obtained a higher average F1 score (0.389 vs. 0.342), although the differences was not statistically significant.

The comparison based on plaque counts showed a different behaviour. While overall MAE was similar between the two methods across the complete dataset (20.52 vs. 21.79 plaques), PyPlaque achieved a lower MAE on the MAYV/CVB3 dataset (2.62 vs. 3.88 plaques). These results indicate that similar counting errors may arise from different instance-level detection quality, highlighting the importance of evaluating both plaque counts and individual plaque localization.

Representative comparisons are shown in Fig. 6. Although both methods operate on identical segmentation masks, the proposed post-processing more effectively separates overlapping plaques, whereas PyPlaque tends to merge adjacent plaques into single detections. Conversely, PyPlaque occasionally produces fewer false-positive detections, illustrating the trade-off between recovering overlapping plaques and suppressing small spurious detections.

**Fig 6.**
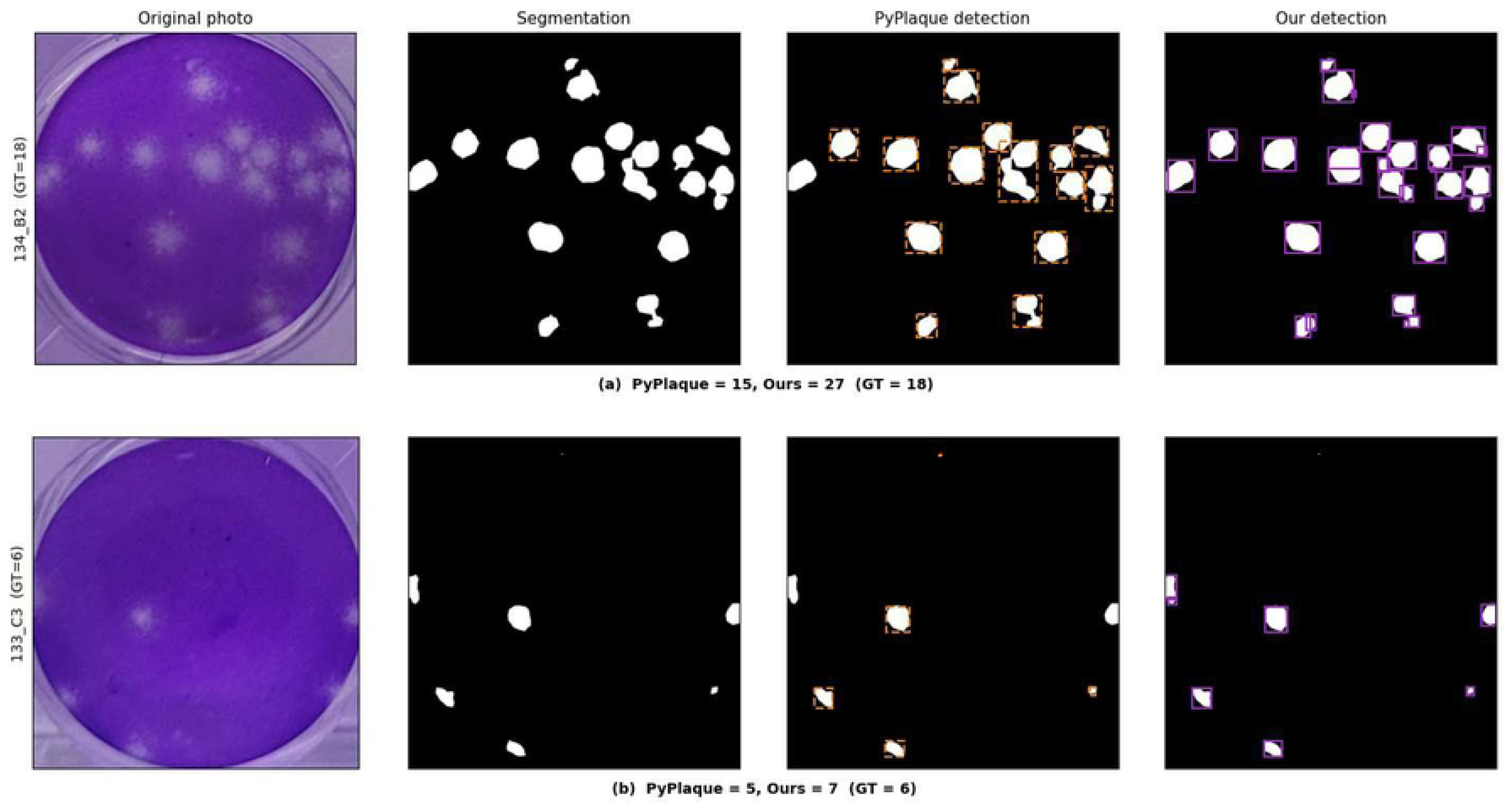
Comparison of plaque detections obtained by applying PyPlaque and our proposed post-processing method to the same predicted segmentation masks in two representative MAYV/CVB3 wells. Columns show the original image, predicted segmentation mask, PyPlaque detections (orange boxes), and detections from our method (purple boxes).

### Plaque density influence

To evaluate the effect of plaque density on automated plaque quantification, test wells were grouped according to their ground-truth (GT) plaque counts. Plaque-detection performance was evaluated using the F1 score, while the mean absolute error (MAE) between predicted and manual plaque counts was used to assess counting accuracy within each density interval (Fig. 7).

**Fig 7.**
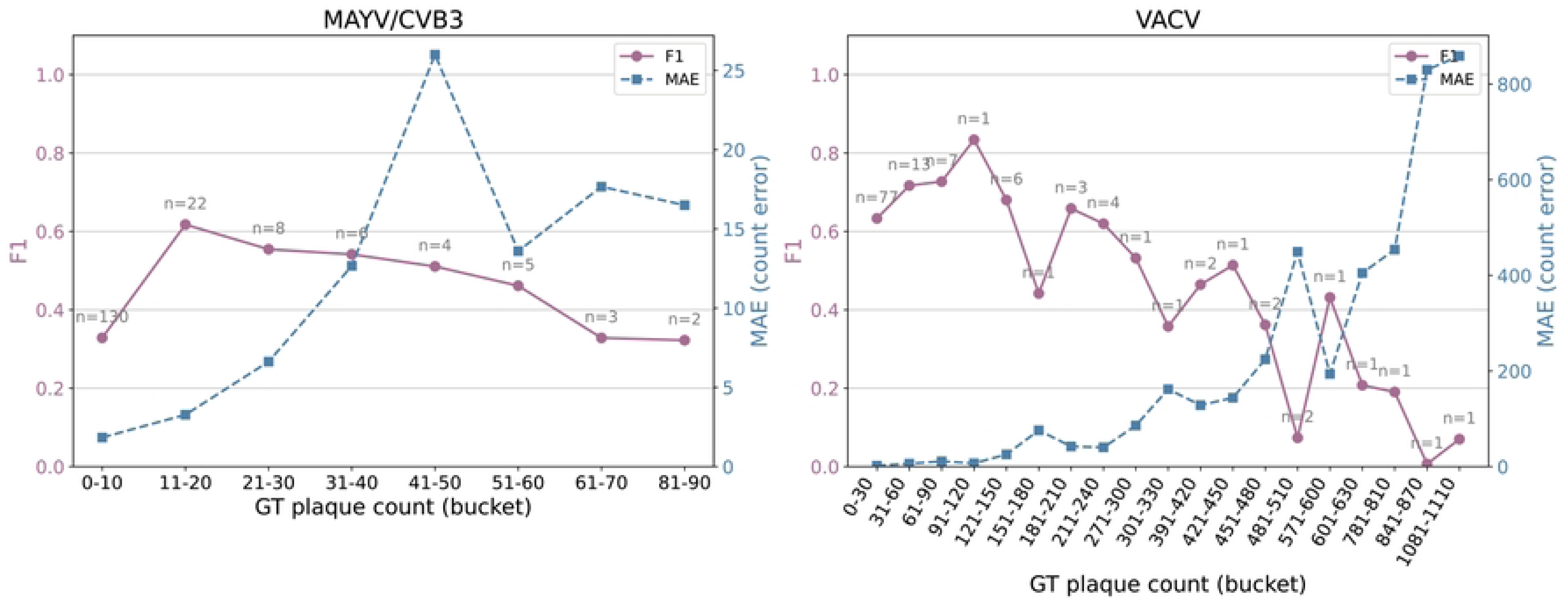
Plaque-detection performance and counting error as a function of ground-truth (GT) plaque count for the MAYV/CVB3 (left) and VACV (right) datasets. Wells were grouped into 10-plaque intervals for MAYV/CVB3 and 30-plaque intervals for VACV. The F1 score (solid line, left y-axis) measures plaque-detection performance, whereas the mean absolute error (MAE; dashed line, right y-axis) represents the mean absolute difference between predicted and manual plaque counts for wells within each interval. Connecting lines illustrate changes in performance with increasing plaque density. Labels above the F1 points indicate the number of wells (*n*) in each interval.

For the MAYV/CVB3 dataset, detection performance was highest for wells containing 11–20 plaques (F1 ≈ 0.62) and gradually decreased as plaque density increased. The lowest-density interval (0–10 plaques) also showed reduced performance despite containing the largest number of evaluated wells (*n* = 130). Counting error showed the opposite behavior overall, increasing with plaque density, although some variability was observed across the higher-density intervals.

A similar but more pronounced density-dependent pattern was observed for the VACV dataset. Detection performance was highest at low-to-intermediate plaque densities and generally declined as plaque density increased, while counting error increased substantially at higher densities. In particular, the highest-density wells showed very low F1 scores together with large counting errors, consistent with reduced counting accuracy under highly confluent conditions. However, the highest-density intervals contained only one or two wells each, limiting the interpretation of performance at these extreme plaque counts.

### Relative counting accuracy and false-positive rate in empty wells

Figure 8 summarizes the relative plaque-counting error of the proposed workflow for wells containing plaques and the frequency of false-positive detections in empty wells.

**Fig 8.**
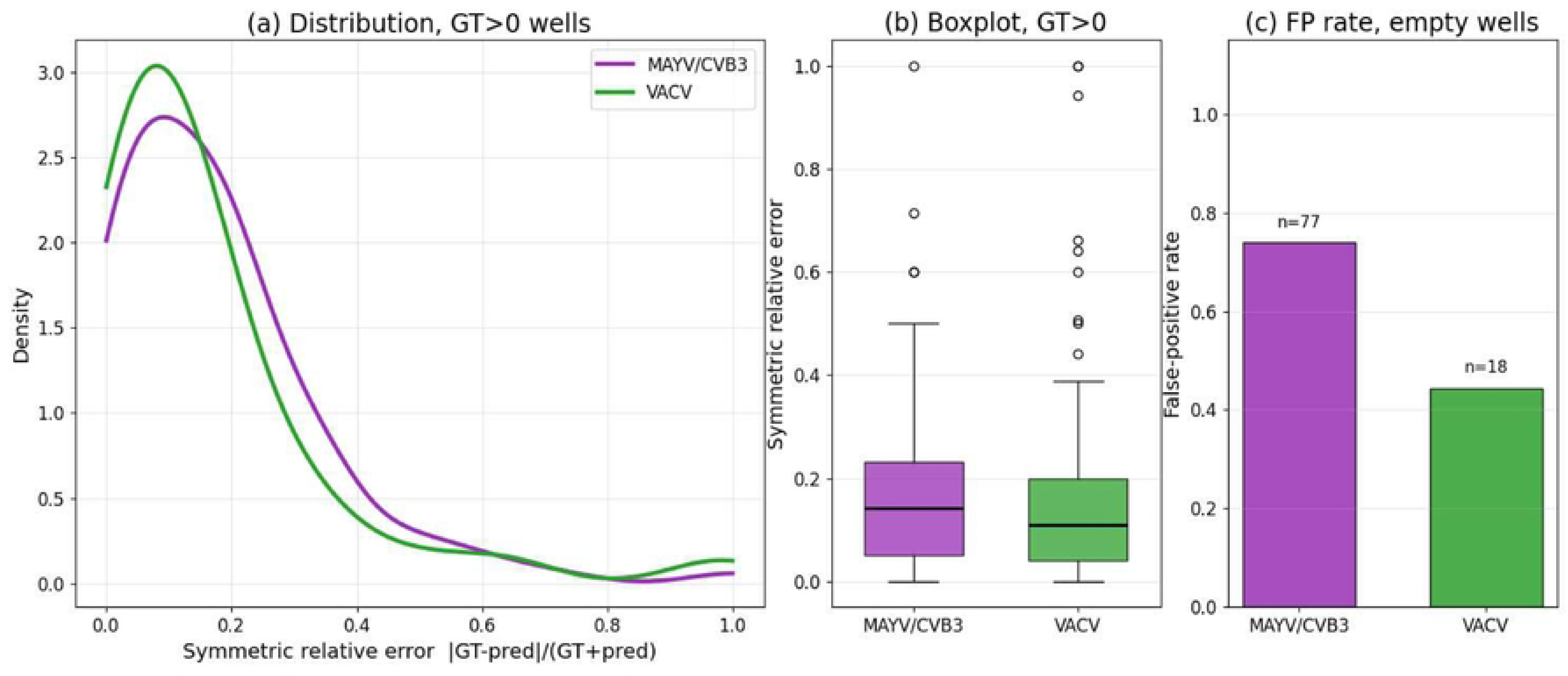
Relative plaque-counting error and false-positive detections of Titra on the MAYV/CVB3 and VACV datasets. (a) Distribution of the symmetric relative counting error for wells containing at least one annotated plaque (GT *>* 0). (b) Boxplot summary of the same distributions. (c) False-positive rate in wells without annotated plaques (GT = 0).

The distributions of symmetric relative counting error were similar for both datasets (Fig. 8a,b), with most wells exhibiting errors below 0.2. Although the median error was slightly higher for MAYV/CVB3 than for VACV (0.14 vs. 0.11), this difference was not statistically significant (Mann–Whitney *U* = 6197, *p* = 0.15), indicating comparable counting accuracy despite the markedly different plaque-density ranges represented by the two datasets.

False-positive detections were more frequent in empty MAYV/CVB3 wells than in VACV wells (0.74 vs. 0.44; Fig. 8c), indicating that plaque-like image structures were more frequently detected under the MAYV/CVB3 imaging conditions.

### Comparison with experts

To place the automated plaque counts in the context of human annotation variability, we compared the proposed workflow with plaque counts provided by four independent experts on two MAYV/CVB3 plates representing different plaque-density profiles (Fig. 9). The middle panels compare the automated counts with each expert individually using scatter plots and Pearson correlation coefficients, while the bottom panels summarize the automated counts together with the expert mean, standard deviation, and individual annotations for each well position.

**Fig 9.**
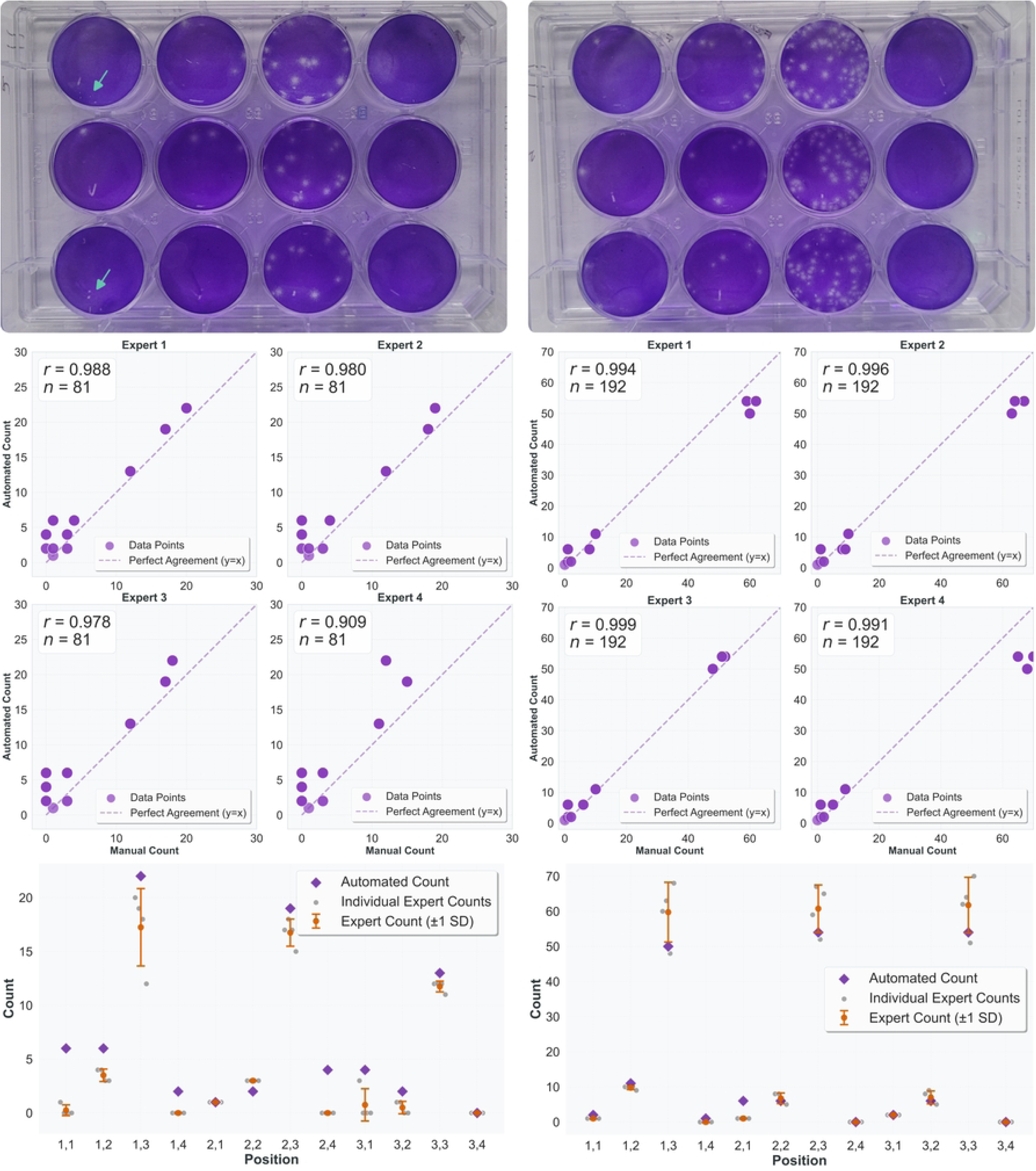
Comparison between automated plaque counts and four independent expert annotations. Top: representative plaque assay plates with examples of small plaque-like regions that were inconsistently annotated by the experts (green arrows). Middle: correlation between automated counts and each expert. Bottom: automated plaque counts compared with the individual expert counts and the expert mean ± SD for each well.

Across both evaluated plates, automated plaque counts showed strong agreement with all four experts. Pearson correlation coefficients ranged from 0.909 to 0.988 for the first plate and from 0.994 to 0.999 for the second plate (Fig. 9), demonstrating high consistency between the proposed workflow and manual annotations.

Most discrepancies between automated and manual counts occurred in wells containing either very small plaque-like structures or high plaque densities. In the first plate, the automated workflow detected several small plaque-like regions (green arrows in Fig. 9) that were not consistently annotated by all experts, leading to higher automated counts in a small number of wells. In the second plate, differences were primarily associated with wells containing larger plaque numbers, where both automated and manual counting became more variable.

The per-well comparison further illustrates that inter-expert variability increased with plaque density. For most wells, automated plaque counts fell within or close to the range of expert annotations, whereas the largest deviations coincided with wells that also exhibited greater disagreement among the experts. These results indicate that many of the remaining differences between automated and manual plaque counts occur in biologically ambiguous cases for which no single expert consensus exists.

### PFU/mL estimation results

To evaluate whether automated plaque counting preserves the final biological measurement, PFU/mL values calculated from automated plaque counts were compared with those obtained from manual annotations for the MAYV/CVB3 dataset (Fig. 10).

**Fig 10.**
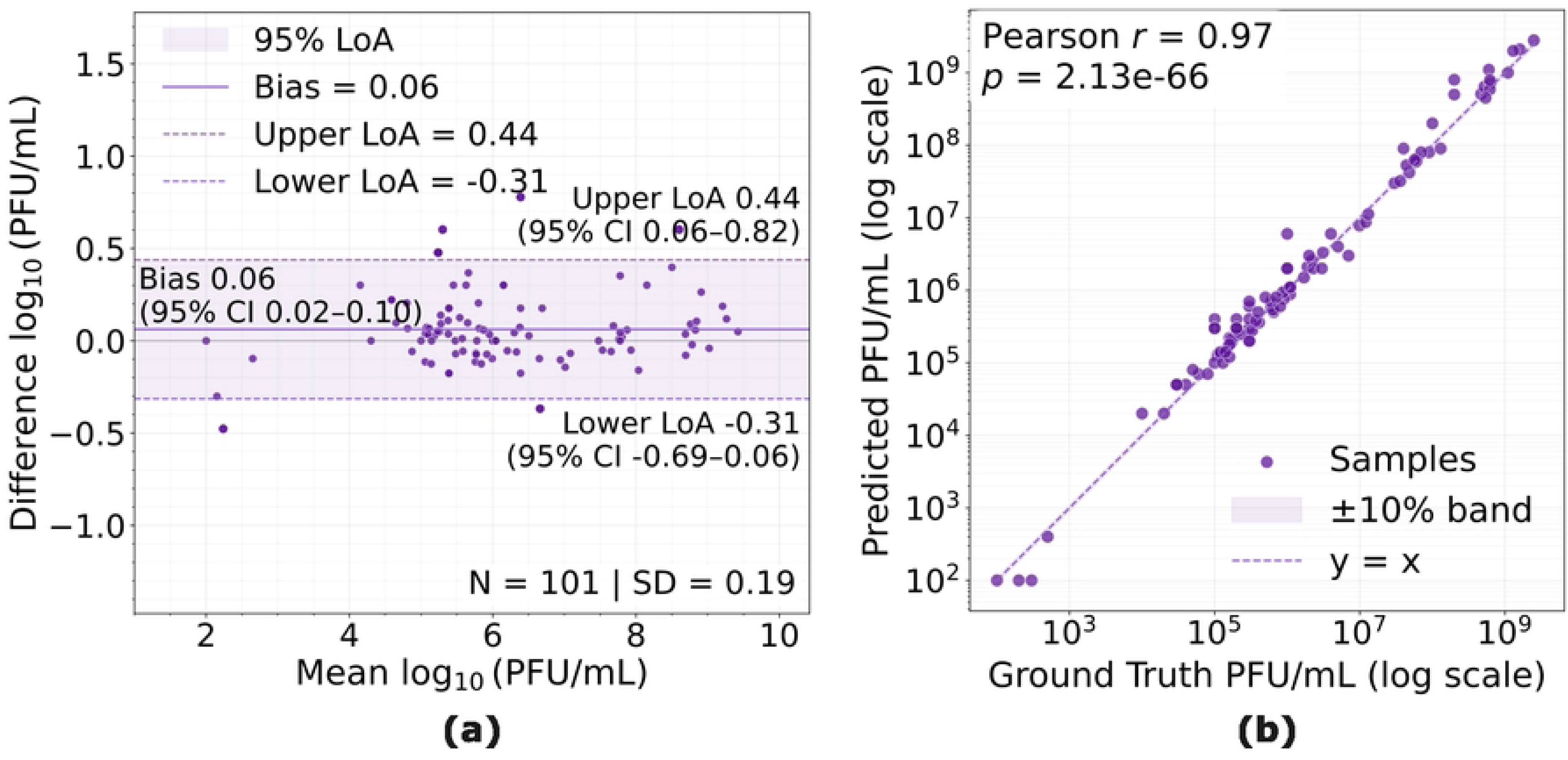
Comparison of PFU/mL values computed for the MAYV/CVB3 dataset using plaque counts obtained from manual annotation and automated plaque detection. a: Bland–Altman plot with agreement between the two PFU/mL measurements. b: Scatter plot comparing PFU/mL values derived from manual and automated plaque segmentations.

The Bland–Altman analysis (Fig. 10a) demonstrated strong agreement between automated and manual PFU/mL estimates. A total of 174 of the 180 evaluated samples fell within the 95% limits of agreement, with a mean difference (bias) of 0.06 PFU/mL and limits of agreement ranging from −0.31 to 0.44. No systematic trend was observed across the range of PFU/mL values.

The corresponding scatter plot (Fig. 10b) showed a strong linear relationship between automated and manual PFU/mL estimates (Pearson *r* = 0.975, *N* = 180), indicating that automated plaque counts produced PFU/mL values that closely matched those obtained through manual plaque enumeration.

### Robustness to image transformations

To evaluate the robustness of the proposed workflow to common image acquisition variations, we assessed well and plaque detection under progressive changes in image brightness and in-plane rotation (Fig. 11).

**Fig 11.**
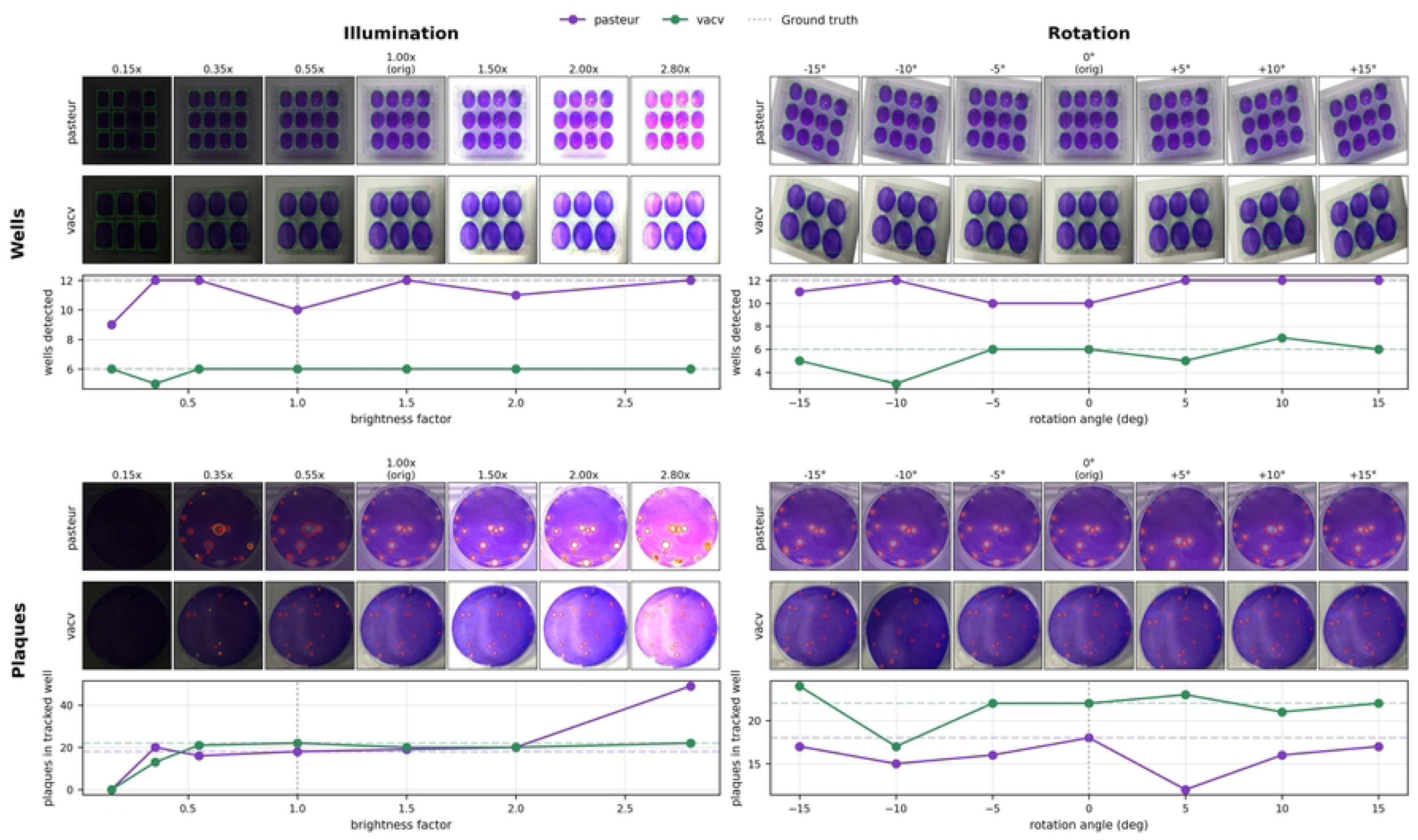
Robustness of well and plaque detection under changes in image brightness and in-plane rotation. Representative transformed images from the MAYV/CVB3 and VACV datasets are shown together with the corresponding well and plaque detections. The plots summarize the number of detected wells and plaques across the evaluated transformations. Horizontal dashed lines indicate the ground-truth counts, and vertical dotted lines indicate the original image (brightness factor 1.0 and rotation 0*^◦^*).

Across both datasets, well detection remained stable over a broad range of brightness levels and rotations, with the correct number of wells detected under most evaluated transformations. Plaque detection exhibited a similar behaviour, with plaque counts remaining close to the corresponding ground-truth values for moderate brightness changes and rotations. Noticeable deviations in plaque counts occurred only under the most extreme transformations, particularly at very low illumination levels, where image contrast was substantially reduced.

Representative transformed images together with the corresponding detection results are shown in Fig. 11.

Overall, these experiments indicate that the proposed workflow maintains stable performance under the moderate acquisition variations typically encountered during routine plaque assay imaging.

## Discussion

In this study, we presented Titra, an end-to-end workflow that automates the complete plaque assay analysis process, from laboratory plate photographs to PFU/mL estimation. Unlike previous approaches, which primarily automate isolated stages such as plaque segmentation or plaque counting, the proposed workflow integrates automatic well detection, plaque segmentation, instance separation, plaque quantification, PFU calculation, and expert review within a single platform. Across three viral species, two plate formats, and heterogeneous acquisition conditions, the workflow produced plaque counts and PFU estimates that closely matched manual analysis while substantially reducing manual intervention. Taken together, these results demonstrate that reliable virus titration does not necessarily require perfect plaque segmentation. Although segmentation performance varied across datasets, downstream plaque counts and PFU estimates remained highly concordant with manual measurements, indicating that the biological endpoint is more robust than pixel-level segmentation metrics alone would suggest.

Rather than treating plaque segmentation as the final objective, Titra considers it one component of a broader workflow whose ultimate goal is reliable virus titration. Existing tools frequently require manual well selection, parameter adjustment, separate PFU calculation, or external software for experiment management. HSD-WBR [27] represents an important advance by incorporating automatic well detection, but it was specifically developed for 6-well plates. In contrast, the proposed workflow automatically analyzes both 6- and 12-well plates, performs plaque quantification and PFU estimation within the same platform, and allows expert review only when necessary. Furthermore, the workflow remained stable under moderate variations in illumination, image rotation, and cropping (Figs. 2 and 11), suggesting that it can tolerate the variability commonly encountered during routine laboratory image acquisition while reducing operator-dependent variability.

The comparison with existing methods highlights that the relative performance of plaque-analysis approaches depends both on the dataset and on the level at which performance is evaluated. For plaque counting, U-Net achieved the lowest error on MAYV/CVB3, followed by Titra, whereas Titra achieved the lowest error on VACV. The corresponding segmentation results showed a similar dataset-dependent behavior: U-Net performed particularly well on MAYV/CVB3, while Titra achieved the strongest Dice, mAP, and detection F1 on VACV. These differences may reflect variations in plaque morphology, density, contrast, and acquisition characteristics between datasets. In particular, the strong performance of Titra on VACV demonstrates its ability to accommodate substantially different plaque morphologies and higher plaque-density conditions. However, evaluation across additional viruses, laboratories, and imaging protocols would be required to establish broader domain generalization.

Importantly, the ranking of methods according to segmentation and detection metrics did not always correspond to their counting performance. This was particularly evident on MAYV/CVB3, where StarDist and HSD-WBR achieved higher Dice and detection F1 scores than Titra but produced larger counting errors. The error analysis provides insight into this apparent discrepancy. Titra generated false-positive detections relatively frequently in empty wells, but these errors typically consisted of only a small number of spurious plaques. StarDist showed the opposite pattern: false-positive failures in empty wells occurred less frequently but could result in large overcounts. These different error profiles are emphasized differently by the evaluation metrics. Macro-averaged detection F1 is strongly affected by the frequency of wells containing detection errors, whereas counting MAE is sensitive to their magnitude. Consequently, occasional severe overcounting can substantially increase MAE without producing an equivalent change in macro-F1, while frequent but small errors can strongly affect macro-F1 while having a comparatively limited effect on the final count. This illustrates why neither metric alone fully characterizes performance for automated plaque quantification.

Pixel-level segmentation metrics provide another complementary perspective. Dice measures spatial agreement between predicted and reference masks and therefore depends on accurate delineation of plaque boundaries. However, plaque borders can be diffuse and low-contrast, making their exact extent difficult to define consistently. A prediction may therefore correctly identify a plaque while differing from the reference annotation around its boundary, reducing Dice without necessarily affecting the resulting plaque count.

Conversely, high pixel-level overlap does not guarantee that touching plaques will be correctly separated into individual instances. This distinction is particularly relevant for plaque assays, where the downstream objective is not precise contour reconstruction but reliable identification and quantification of individual plaques. The comparison with PyPlaque further supports the importance of this intermediate step, as different strategies for separating overlapping plaques can produce different instance-level and counting outcomes even when operating on the same segmentation masks.

Together, these findings indicate that automated plaque-assay methods should not be evaluated using a single segmentation or detection metric. Dice and mAP characterize segmentation quality, detection F1 captures instance-level identification, and counting error directly measures the downstream quantitative task.

Their combination provides a more complete characterization of both overall performance and method-specific failure modes. For Titra, this analysis also identifies a concrete opportunity for improvement: reducing the small false-positive detections observed in empty or low-density wells, for example through size-aware filtering or additional confidence-based rejection, could improve instance-level performance without altering the broader workflow.

Analysis of the remaining errors consistently identified plaque density as an important factor limiting automated plaque quantification (Fig. 7). As plaque density increased, overlapping plaques progressively obscured individual boundaries, reducing instance separation and leading to undercounting. This behavior was consistently observed across the qualitative examples, density-stratified analysis, and Bland–Altman plots (Fig. 4). The effect was particularly pronounced in VACV, where decreasing detection F1 at high plaque densities was accompanied by increasing counting errors, consistent with the increasing difficulty of separating highly overlapping and confluent plaques. In contrast, MAYV/CVB3 covered a substantially lower plaque-density range, preventing us from determining whether a similar degradation would occur at comparable densities. Moreover, the highest-density VACV intervals contained only a small number of wells, and performance under these extreme conditions should therefore be interpreted cautiously. Together, these results identify highly confluent wells as an important scenario for further methodological improvement.

Comparison with four independent experts provided additional context for interpreting automated performance (Fig. 9). The largest discrepancies between automated and manual counts occurred for small plaque-like structures and highly ambiguous regions, which also showed the greatest disagreement among the experts themselves. Consequently, part of the observed disagreement reflects the intrinsic uncertainty of plaque annotation rather than algorithmic error alone. This observation reinforces the value of incorporating expert review within the workflow, allowing ambiguous cases to be resolved rapidly while benefiting from automated analysis for the majority of samples.

Although the robustness experiments demonstrated stable performance under moderate variations in illumination, image rotation, and cropping, and evaluation across Mayaro virus, Coxsackievirus B3, and vaccinia virus encompassed different plaque morphologies, plaque densities, and acquisition conditions, these results should not be interpreted as evidence of broad domain generalization. Rather, they demonstrate that the proposed workflow can operate successfully beyond a single experimental setting. Additional validation across a larger number of viral species, laboratories, imaging devices, and experimental protocols will be necessary before broader generalization can be claimed.

Finally, the modular architecture of Titra provides opportunities beyond conventional plaque assays.

Because the workflow separates well localization, plaque analysis, and biological quantification into independent components, it could be adapted to related assays such as plaque reduction neutralization tests (PRNT) and potentially *TCID*_50_ assays following task-specific validation. Future studies should also focus on improving analysis of highly confluent plaques, incorporating biologically informed priors such as adaptive plaque-size filtering, and leveraging expert corrections collected through the review interface to continually refine the underlying models. Together, these directions position Titra as a flexible human-in-the-loop platform for automated plaque assay analysis that can continue evolving as additional datasets, laboratory protocols, and foundation models become available.

## Conclusion

We developed an end-to-end workflow that automatically detects assay wells, segments and counts plaques, and computes PFU/mL estimates directly from laboratory plaque assay images. Evaluation on images from Mayaro virus, Coxsackievirus B3, and vaccinia virus demonstrated that the proposed workflow provides reliable plaque quantification across the evaluated datasets while remaining robust to different plate formats and acquisition conditions. Although plaque overlap in highly confluent wells remains a challenge, automated plaque counts and PFU/mL estimates showed strong agreement with manual analysis, supporting the suitability of the workflow for routine laboratory use.

By combining automated image analysis with expert review through the *Titra* platform, the proposed approach improves the reproducibility and scalability of plaque assay quantification while preserving expert oversight for biologically ambiguous cases. Future studies will focus on validating the workflow across a broader range of viral species and laboratory settings, as well as extending the modular architecture to additional virological assays.

## Acknowledgements

We thank Mercedes Paz, Natalia Echeverría, Álvaro Fajardo, Paula Perbolianachis, and Juan Gandioli for their participation in testing Titra and for their valuable feedback during its evaluation.

## Author contributions statement

All authors contributed to the conceptualization and design of the solution. E.M. and J.I.O. designed, implemented and validated deep learning models. L.L.L., M.V., A.E.V., J.R., I.M. designed and implemented the web application. A.C., S.R., I.F. and J.H. conducted the viral assay experiments and provided both image data and their annotations. E.M., A.C., P.M., G.M. and J.I.O. analysed the results. E.M. and J.I.O. wrote the manuscript. All authors reviewed the manuscript.

## Additional information

### Data availability

The VACVPlaque benchmark images used in this study are openly available on RODARE [12] under a CC-BY 4.0 license. The remaining primary images were collected at Institut Pasteur de Montevideo under institutional agreements; these raw images will be published by the time of publication. Derived annotations (well and plaque masks) and aggregate titration results required to reproduce our analyses will be publicly released upon acceptance of this manuscript.

### Code availability

The Titra platform and the end-to-end analysis pipeline will be released as open-source software at github.com/arionkoder/titra-ai upon acceptance of this paper.

